# Cryo-EM structure of a blue-shifted channelrhodopsin from *Klebsormidium nitens*

**DOI:** 10.1101/2024.01.21.576531

**Authors:** Yuzhu Z. Wang, Koki Natsume, Tatsuki Tanaka, Shoko Hososhima, Rintaro Tashiro, Fumiya K. Sano, Hiroaki Akasaka, Satoshi P. Tsunoda, Wataru Shihoya, Hideki Kandori, Osamu Nureki

**Author notes:** To whom correspondence should be addressed. (W.S.), Hideki Kandori (H.K.), and (O.N.).

## Abstract

Channelrhodopsins (ChRs) are light-gated ion channels and invaluable tools for optogenetic applications. Recent developments in multicolor optogenetics, in which different neurons are controlled by multiple colors of light simultaneously, have increased the demand for ChR mutants with more distant absorption wavelengths. Here we report the 2.9 Å-resolution cryo-electron microscopy structure of a ChR from *Klebsormidium nitens* (KnChR), which is one of the most blue-shifted ChRs. The structure elucidates the 6-*s-cis* configuration of the retinal chromophore, indicating its contribution to a distinctive blue shift in action spectra. The unique architecture of the C-terminal region reveals its role in the allosteric modulation of channel kinetics, enhancing our understanding of its functional dynamics. Based on the structure-guided design, we developed mutants with blue-shifted action spectra. Finally, we confirm that UV or deep-blue light can activate KnChR-transfected precultured neurons, expanding its utility in optogenetic applications. Our findings contribute valuable insights to advance optogenetic tools and enable refined capabilities in neuroscience experiments.

## Main text Introduction

Channelrhodopsins (ChRs) are light-gated ion channels in the eyespot of green algae. Channelrhodopsin-1 and -2, derived from *Chlamydomonas reinhardtii* (CrChR1 and CrChR2), were the first ChRs to be discovered and characterized, and the latter has been extensively utilized for optogenetic applications^1^. These proteins conduct cations such as H^+^, Na^+^, K^+^, and Ca^2+^ ^2^. Cation channelrhodopsins with sequence homology to ChR2 are denominated as chlorophyte CCRs^3^. X-ray structures of a ChR chimera of C1C2 and CrChR2 have unveiled intricate details regarding the molecular architecture^4^, shedding light on the photoactivation and ion conduction pathway. ChRs have been employed to elicit action potentials in light-insensitive cells and tissues with unparalleled spatiotemporal precision, thereby launching a nascent research discipline, optogenetics^3^. ChR variants have been iteratively engineered to improve their functionalities^5^. Furthermore, genome mining has expedited the discovery of novel ChRs and their applications for optogenetics.

The mining and engineering of ChRs has facilitated multicolor optogenetics^6–17^. This technology leverages red and blue opsins to concurrently regulate two distinct neurons. It is particularly useful in synaptic communication, as an invaluable asset for probing the intricacies of complex neural circuits. Among the red opsins, ChrimsonR^6^ is commonly employed, and the recently discovered ChRmine^18^ harbors the potential to surpass ChrimsonR. In the realm of blue opsins, popular mutant variants of CrChR2 have been adopted^5^ alongside recently identified ChRs such as sdChR and Chronos^6^. A rational approach to creating color variants has been applied, producing the blue-shifted color variant C1C2GA^19^. Nevertheless, many challenges remain for the application of blue opsins in multicolor optogenetics. To minimize crosstalk with red opsins, the activation of blue opsins with blue light (405 nm) is imperative^9,17^, as this wavelength does not stimulate red opsins. This limitation may generate inadequate excitation by low levels of blue opsin expression, consequently impeding the utility of multicolor optogenetics. In light of these considerations, a blue opsin with shorter wavelength excitation is keenly desired for use in multicolor optogenetics.

Among the most pronounced blue-shifted ChRs is a recently discovered ChR derived from the filamentous terrestrial alga *Klebsormidium nitens*, designated as KnChR^20^. KnChR comprises a 7-transmembrane rhodopsin domain that contains a channel pore, followed by a C-terminal moiety that encodes a peptidoglycan binding domain known as FimV. KnChR exhibits higher Ca^2+^ permeability and about 10-fold higher metal cation permeability relative to H^+^, as compared with CrChR2. KnChR exhibits maximal sensitivities at 430 and 460 nm, with the former making KnChR one of the most notable blue-shifted ChRs to date, with the potential to serve as a pioneering framework for exploring the molecular mechanisms governing ChR color tuning. Notably, the rate of channel closure is impacted by the C-terminal moiety, and the truncation of this moiety led to over 10-fold prolongation of the channel’s open lifetime.

Two pivotal arginine residues, R287 and R291, play a crucial role in modulating the photocurrent kinetics. However, little is known about how the C-terminal residues influence the channel activity and the mechanisms governing short wavelength excitation, as well as their optogenetic utility.

## Results

### Structure determination

To examine mechanisms of the blue-shifted excitation and channel modulation, we performed a cryo-electron microscopy (cryo-EM) single-particle analysis of KnChR. First, we expressed KnChR, containing 697 amino acids, in insect cells and purified it in detergent micelles (Supplementary Fig. 1a, b). However, during the cryo-EM analysis, only the top view was visible in the 2D class averages (Supplementary Fig. 1c). This limitation was attributed to either the orientation bias or the deleterious effects of the micelles. Consequently, we reconstituted purified KnChR into MSP2N2 nanodiscs (Supplementary Fig. 2a) and performed the cryo-EM structural analysis. The nanodisc reconstitution eliminated the orientation bias, and the side view became visible (Supplementary Fig. 2b). Finally, we determined the structure of KnChR in the ground state at a nominal resolution of 2.9 Å (Fig. 1a, Table 1, Supplementary Fig. 2b).

**Fig. 1.**
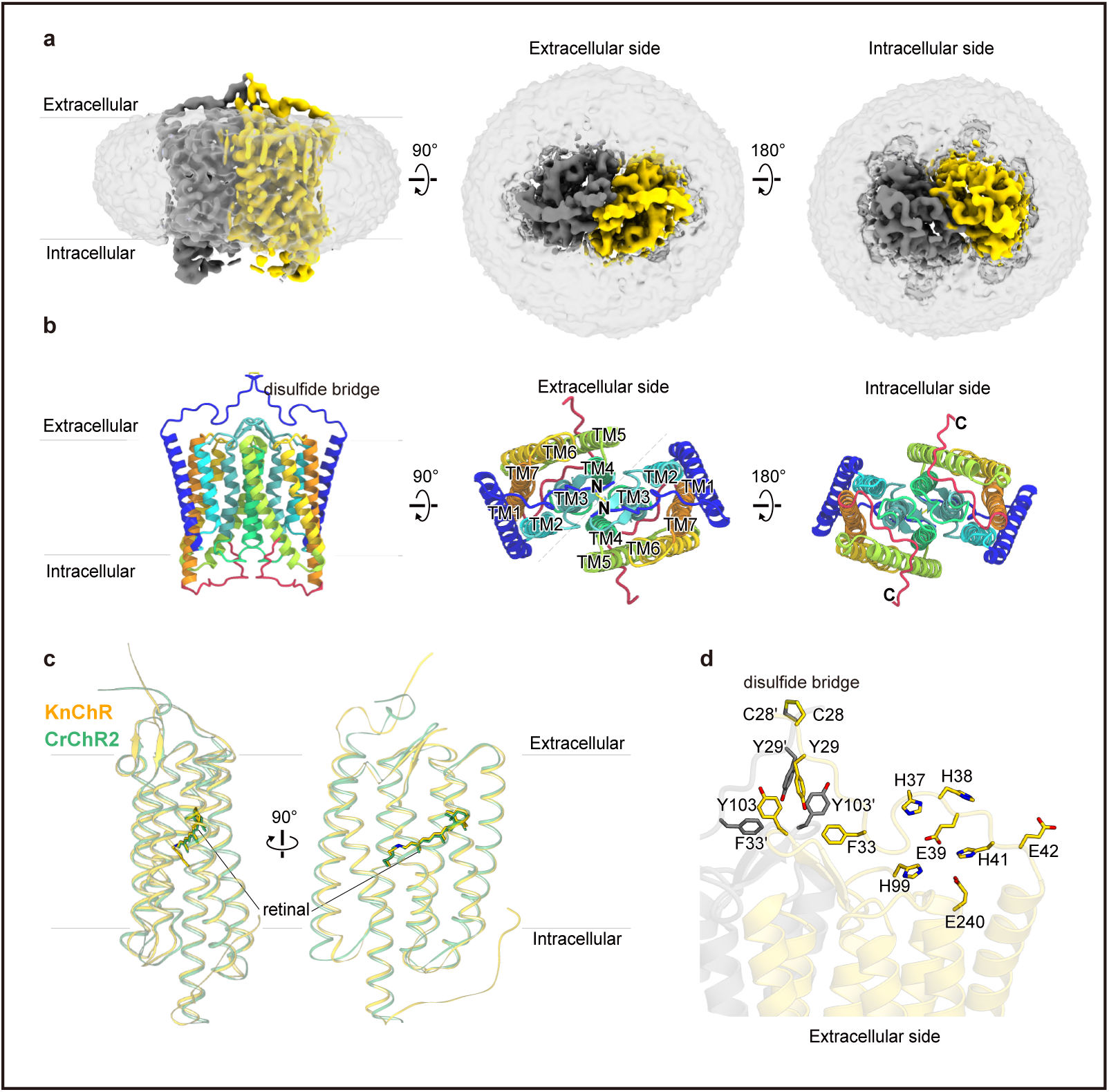
Overall structure of the KnChR dimer. **a**, Overall structure of the nanodisc-reconstituted KnChR dimer. The cryo-EM density maps are shown with the two protomers in different colors. **b**, Ribbon representation of KnChR. Each KnChR protomer is depicted in a cartoon model, with the N-terminal region in blue and the C-terminal region in red. 6-*s-cis* retinal (stick model) is embedded within the 7TMs. The intermolecular disulfide bridge, formed by C28, is shown as a yellow stick. **c**, Superimposition of KnChR and CrChR2 (PDB ID: 6EID). **d**, Characteristic interactions in the N-terminal region.

**Table 1.** Cryo-EM data collection, refinement, and validation statistics.

| **Data collection** | KnChR |
| --- | --- |
| Microscope | Titan Krios (Thermo Fisher Scientific) |
| Voltage (keV) | 300 |
| Electron exposure (e<sup>-</sup>/Å<sup>2</sup>) | 49.263 |
| Detector | Gatan K3 Summit camera (Gatan) |
| Magnification | ×105,000 |
| Defocus range (μm) | −0.6–1.6 |
| Pixel size (Å/pix) | 0.83 |
| Number of movies | 8,554 |
| Symmetry | C2 |
| Picked particles | 3,102,701 |
| Final particles | 87,244 |
| Map resolution (Å) | 2.91 |
| FSC threshold | 0.143 |
| **Model refinement** |  |
| Atoms | 2164 |
| **R.m.s. deviations from ideal** |  |
| Bond lengths (Å) | 0.003 |
| Bond angles (°) | 0.553 |
| Validation |  |
| Clashscore | 4.15 |
| Rotamers (%) | 0.00 |
| **Ramachandran plot** |  |
| Favored (%) | 98.48 |
| Allowed (%) | 1.52 |
| Outlier (%) | 0 |

Like other chlorophyte CCRs, KnChR forms a dimer and comprises transmembrane helices ^21^(TMs) 1–7, as well as N- and C-terminal regions (Fig. 1b). We successfully modeled residues 27 to 291. Notably, the C-terminal FimV domain was disordered (Supplementary Fig. 1a, 2b), indicating its inherent structural flexibility. This observation aligns with the previous report^20^ that the FimV truncation does not alter the channel activity. The retinal chromophore is covalently bound to K254, forming the retinal Schiff base (RSB)^2^. The local resolution surrounding the central retinal is approximately 2.6 Å (Supplementary Fig. 2b), while the N- and C-terminal regions exhibit a comparatively lower resolution of 3.8 Å. Nevertheless, this resolution is adequate for modeling protein residues. The rhodopsin domain of KnChR has 34.5% identity and 69.3% homology with CrChR2 and superimposed well on CrChR2^4^ (PDB ID: 6EID), with a root mean square deviation of 1.15 Å^4^ (Fig. 1c). The N-terminal region extends to the center of the dimer interface, and C28 of each protomer forms a disulfide bond with its counterpart. These structural features represent the typical architecture of chlorophyte CCRs^2^ (Supplementary Fig. 3a–e)^4,19,22,23^.

Remarkably, in contrast to other ChRs^22,23^, the N-terminal region adopts an elongated conformation and does not contain any α-helices (Supplementary Fig. 3a–e), and is stabilized by extensive electrostatic interactions between histidine residues (H37, H38, H41, and H99) and two glutamate residues (E39 and E240) (Fig. 1d). Moreover, aromatic residue clusters constitute the dimer interface under the C28-C28 disulfide bond. These structural features of the N-terminal region are unique to KnChR.

### Mechanism of blue-shifted excitation

We first attempted to model all-*trans* retinal (ATR) into the density (Fig. 2a), but encountered difficulty in confidently fitting the β-ionone ring. Alternatively, 6-*s-cis* retinal was clearly fitted into the density (Fig. 2b, c, Supplementary Fig. 2b). In ATR, the β-ionone ring lies in the same plane as the polyene chain, due to the fixed planar conformation of the C6–C7 bond (Fig. 2d). Conversely, in 6-*s-cis* retinal, the β-ionone ring undergoes a rotation of approximately 140°, enforced by torsion around the C6–C7 bond, thus deviating from planarity^24^ (Fig. 2c). This rotation leads to a contraction in π-conjugation, ultimately inducing blue shifts in absorption spectra^2,19,25^. Thus, the observed 6-*s-cis* retinal could account for the blue-shifted excitation of KnChR.

**Fig. 2.**
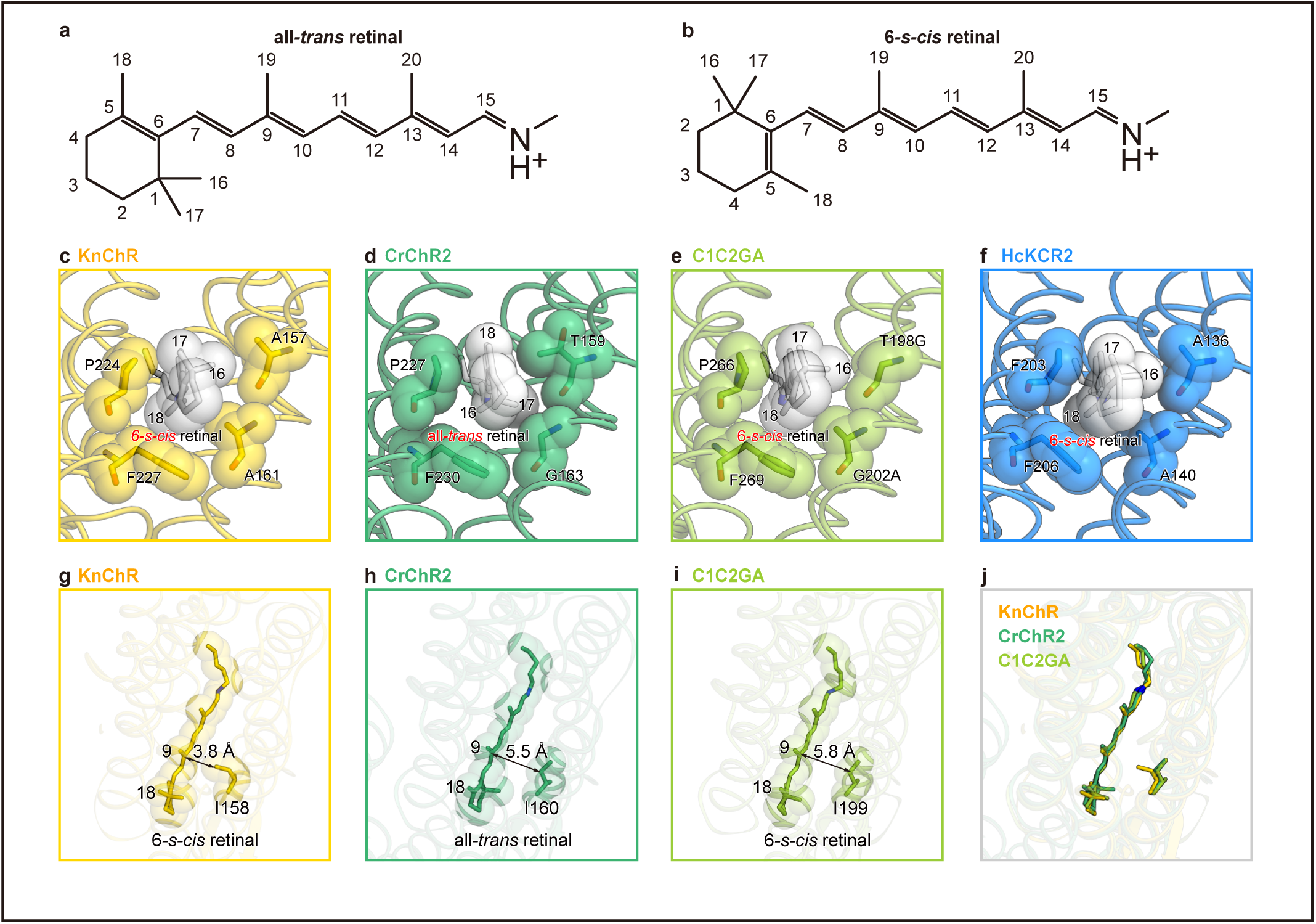
6-*s-cis* configuration. **a**, **b**, Chemical structure of the chromophore, RSB, and the atom numbering of ATR (a) and 6-*s-cis* retinal (**b**). **c–f,** Close-up views of the chromophore binding pocket around the β-ionone ring. Surrounding residues are shown as sticks with transparent cpk models. **g,** Superimposition of KnChR, CrChR2 (PDB ID: 6EID), and C1C2GA (PDB ID:4YZI), focused on the interaction between retinal and I158 in KnChR. **h–j**, Distances between retinal and I158 in KnChR (**h**), I160 in CrChR2 (PDB ID: 6EID) (**i**), and C1C2GA (PDB ID: 4YZI) (**j**). Double-ended arrows indicate the distances.

Notably, a 6-*s-cis* retinal configuration is also observed in HcKCR2 (PDB ID: 8H87)^23^ and the engineered ChR mutant C1C2GA (PDB ID:4YZI)^19^, which exhibit blue-shifted excitations compared to HcKCR1 and C1C2^21^, respectively (Fig. 2e, f). The conversion is induced by the A136/A140 mutation in HcKCR2 and the T198G/G202A mutation in C1C2GA, in agreement with the “Non-G rule^22^”, which is characterized by β-ionone ring rotation owing to steric hindrance. Correspondingly, the homologous residues A157/A161 in KnChR enable the 6-*s-cis* configuration through a mechanism similar to that of HcKCR2.

Upon comparing the other residues that constitute the retinal binding site (Supplementary Fig. 4a, b), I158 in KnChR exhibited a unique structural feature (Fig. 2g, h), although it is highly conserved among the chlorophyte CCRs. In CrChR2, the distance between the homologous I160 and the retinal is 5.5 Å (Fig. 2h)^4^, suggesting the absence of a direct interaction. This holds true for C1C2GA with the 6-*s-cis* retinal configuration (Fig. 2i)^19^. However, in knChR, the I158 side chain diverges more toward the 6-*s-cis* retinal (Fig. 2g), resulting in a distance of 3.8 Å to C9 of the retinal, indicative of a direct interaction (Fig. 2j). While the underlying cause of this feature remains elusive, it may be attributed to structural disparities specific to KnChR. This additional van der Waals interaction with the retinal is hypothesized to elevate the energy level of the excited state, thereby creating a larger energy gap between the ground and excited states, and consequently inducing the observed blue shift.

### Ion pathway

In chlorophyte CCRs, the ion pathways are formed by TM2, 3, 6, and 7 and are constrained by three constriction sites that limit ion permeation: the intracellular (ICS), central part of the membrane (CCS), and extracellular (ECS) constriction sites. The ion pathway in KnChR adheres to this pattern (Fig. 3a), except for the ECS. The extracellular volume (EV) in KnChR is not divided by the ECS, in contrast to CrChR2. In CrChR2^4^, Q117, R120, and H249 primarily constitute the ECS, dividing the extracellular volume (EV) into EV1 and EV2 (Fig. 3b). In KnChR, Q117 is replaced with P115, resulting in a unified EV (Fig. 3a). The single EV is similarly observed in C1C2, indicating the ion pathway for the extracellular space. One of the prominent differences between the structures of KnChR and CrChR2 is the interaction of K90/E94 (K93/E97 in CrChR2). In KnChR, the distance between these residues is 2.62 Å, and they form an electrostatic interaction. However, K93 and E97 are 3.27 Å apart in CrChR2. The KnChR E94 mutation retarded the on- and off-kinetics in the patch clamp measurement (Fig. 3c), whereas minimal kinetic effects were reported for the ChR2 E97 mutation^26^. These results highlight the distinct role of KnChR E94 in the channel gating. At the CCS part, ChR2 forms a characteristic hydrogen bonding network with E90, K93, E123 and D253, whereas only Y91 interacts with D250^4^. In the central part of KnChR, two carboxylates E121 and D250 serve as counterions, thereby stabilizing the positive charge of the RSB (Fig. 3b), as in other chlorophyte CCRs. In CrChR2, E90, one of the key determinants of H^+^ selectivity, forms water-mediated hydrogen bond interactions with E123 and D253, constituting the CCS^4^. In KnChR, the side chain of the homologous residue E87 diverges from the counterions and engages in direct and water-mediated hydrogen bonds with N255 (Fig. 3b). The KnChR E87Q mutant exhibits slower channel-closure kinetics in the patch-clamp measurement (Fig. 3c), suggesting that E87 is responsible for the reprotonation of the RSB.

**Fig. 3.**
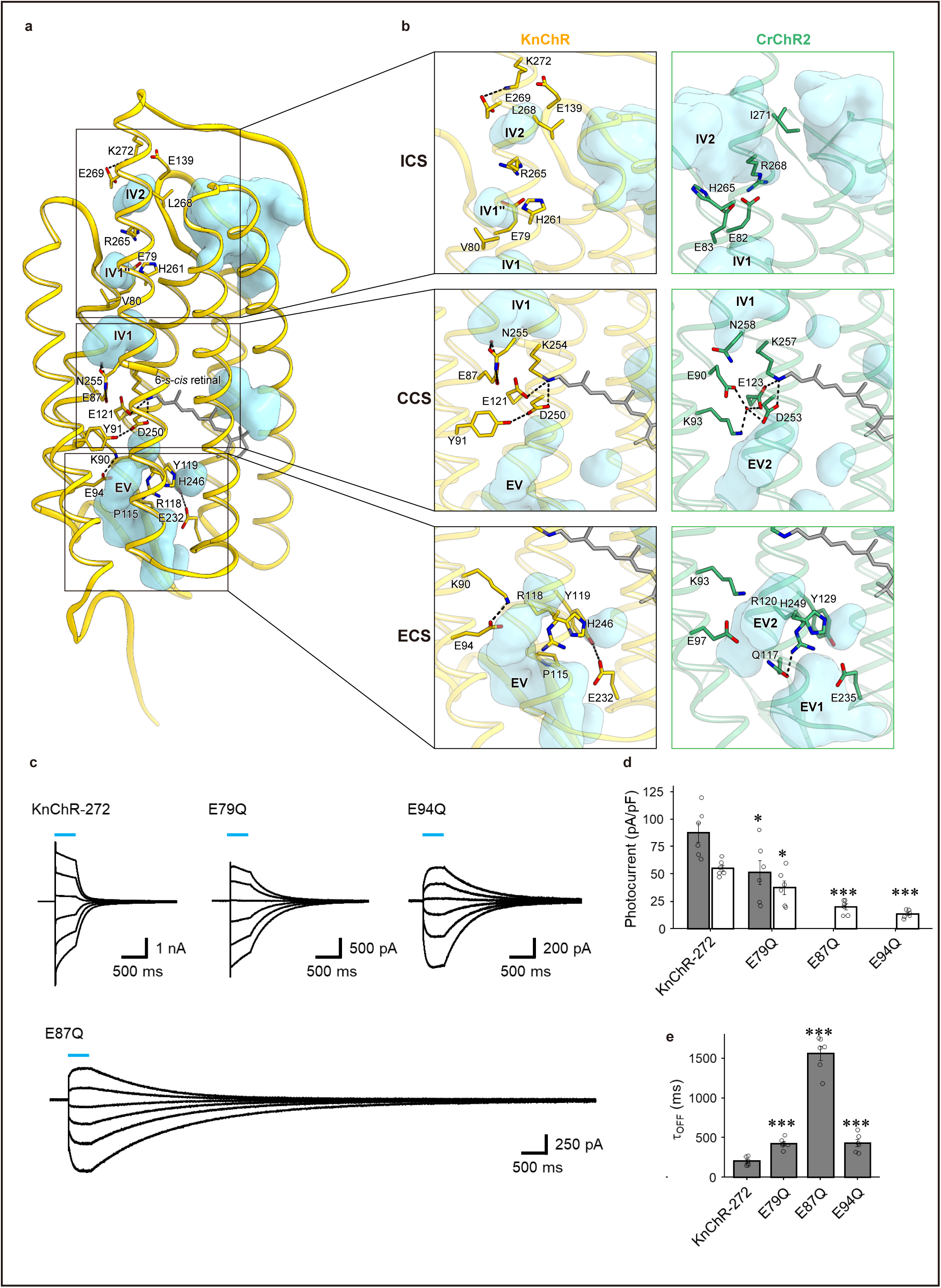
Ion pores of KnChR and CrChR2. **a,** Water accessible cavities are illustrated in the KnChR structure, with the putative ion pathway indicated by an arrow. The three constriction sites for the inner, central, and extracellular gates are enclosed within squares. **b**, Comparison of the constriction sites of KnChR (left panels) and CrChR2 (PDB ID: 6EID) (right panels), for the inner (upper panels), central (middle panels), and outer (lower panels) gates. The constituent residues are shown as sticks, and the TM helix number is indicated on each helix. Black dashed lines indicate hydrogen bonds. c, Representative photocurrent traces of KnChR-272 and mutants. The cells were illuminated with light (λ = 470 nm) during the time indicated by blue bars. The membrane voltage was clamped from −60 to +40 mV for every 20-mV step. **d,** Comparison of photocurrent amplitudes of KnChR-272 and mutants at −60 mV. In the bar graph, gray bars indicate the amplitude from the peak photocurrent (Ip), and open bars indicate the amplitude from the steady-state photocurrent (Iss). n = 6 cells. e, The channel-closing kinetics of KnChR-272 and mutants after illumination cessation (τ-off) at − 60 mV. n = 6 cells.

On the intracellular side of KnChR, E79, H261, and R265 form the ICS (Fig. 3b), dividing the intracellular volume (IV) into two discrete parts (IV1 and IV2) as in CrChR2^4^. IV1 is further divided into IV1 and IV1” by E79 and V80. The E79Q mutation slightly retarded the off-kinetics in the patch-clamp measurement (Fig. 3c), indicating that this residue affects the channel gating. IV2 is smaller in KnChR compared to CrChR2^4^ because of the electrostatic interaction above IV2, featuring E139, E269, and K272. Thus, the differences in ICS, CCS, and ECS between KnChR and CrChR2 (Fig. 3a, b) are likely to contribute to the distinct ion permeabilities of these channels.

### Insight into the C-terminal region

We illuminated the density corresponding to the C-terminal region modulates the channel activity (Supplementary Fig. 2b), and modeled the residues up to R291 (Fig. 4a, b). The C-terminal region extends clockwise from TM7 to TM5, forming extensive interactions with TM5, a unique structural feature in ChRs (Supplementary Fig. 3f-j)^4,19,22,23^. Among the C-terminal residues, R287 exhibits the most significant impact on channel modulation, since the R287A mutation prolongs the lifetime of the channel-opening and increases the current amplitude^20^. In the cryo-EM structure, R287 forms an electrostatic interaction with E196 and a polar interaction with Q192 (Fig. 4c). Thus, we measured the photocurrents of the R287E and E196R mutants, in which the electrical charge is inverted (Fig. 4d). The photocurrent amplitude of the R287E mutant was elevated by ∼220%, and the kinetics of channel-closing (*τ*_off_) slowed down from 25 ms to 160 ms, confirming the importance of the residue for channel gating (Fig. 4e and 4f). By contrast, only a small effect on *τ*_off_ was observed in the E196R mutant, although the current amplitude was increased (Fig. 4d–f). These results indicate that not only the electrostatic interaction but also a polar interaction could be responsible for the channel modulation.

**Fig. 4.**
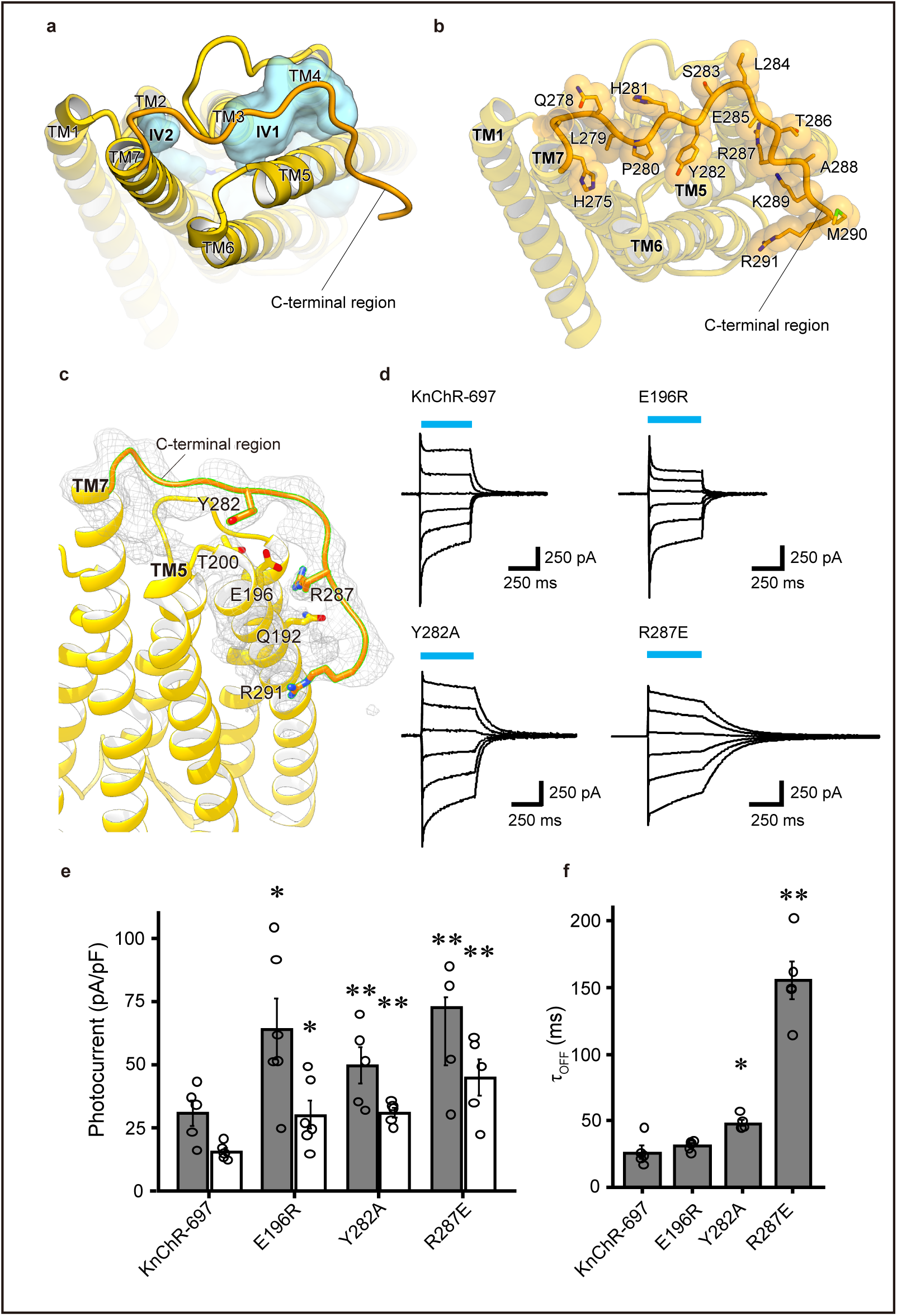
Intracellular face of KnChR. **a,** Water accessible cavities, viewed from the intracellular side. The C-terminal region is highlighted. **b**, The residues in the C-terminal region are shown as sticks with transparent CPK models. **c**, Close-up view of Y282, R287, and R291 with a transparent unsharpened cryo-EM map. **d**, Representative photocurrent traces of KnChR-697 and mutants. The cells were illuminated (λ = 470 nm) during the time indicated by blue bars. The membrane voltage was clamped from −60 to +40 mV for every 20-mV step. **e**, Comparison of photocurrent amplitudes of KnChR-697 and mutants at −60 mV. In the bar graph, gray bars indicate the amplitude from the peak photocurrent (Ip), and open bars indicate the amplitude from the steady-state photocurrent (Iss). n = 5 to 6 cells. **f**, The channel-closing kinetics of KnChR-697 and mutants after illumination cessation (τ-off) at −60 mV. n = 5 to 6 cells.

Moreover, Y282 forms van del Waals interactions with the main and side chains of T200, which are clearly visible in the cryo-EM map(Fig. 4c). The Y282A mutant exhibited a relatively small but a significant change in *τ*_off_. These interactions between the C-terminus and TM5 would be involved in channel modulation.

The C-terminal region faces another spacious IV, rather than IV2 (Fig. 3a, 4a). This voluminous IV is also present in CrChR2, but is not involved in the ion pathway (Fig. 3b)^4^. This observation implies that it exerts allosteric modulation, potentially curtailing conformational shifts and facilitating rapid inactivation in response to light (Fig. 4c). This interaction between the C-terminal region with TM5 distinguishes KnChR from other ChRs, and thus represents a unique structural feature (Supplementary Fig. 3f–j) ^4,19,22,23^.

It should be noted that R291 poorly interacts with the other part of KnChR (Fig. 4c), while the R291A mutation prolonged the lifetime of the channel-opening and increased the current amplitude. The R291 residue is exposed to the membrane environment, and its interactions with the phosphate groups of the lipid bilayer may anchor the C-terminal region.

### Blueshift color-tuning and Optogenetic characterization

ChRs with shorter wavelength excitation are optimal for multicolor optogenetics with dual light applications. Consequently, we explored blue-shifted mutants of KnChR (Fig. 5a, Supplementary Fig. 5a-c). Initially, we focused on A157/A161, which are responsible for the 6-*s-cis* configuration (Fig. 5b). We surmised that the A157G mutant, which mimics the T198G/G202A mutation in C1C2GA, would further stabilize the 6-*s-cis* configuration. Consistently, the A157G mutation (KnChortRG) showed a 40-nm blue-shifted action spectrum (λmax=410 nm) in the patch-clamp recording, as compared with that of the wild-type (λmax=450 nm) (Fig. 5a, c). Subsequently, we focused on the L/Q switch, which is known for its role in color-tuning^27^. For the proteorhodopsin group, the presence of leucine and glutamine at position 105, referred to as the L/Q switch, regulates the spectra between the green- and blue-absorbing types, respectively. The homologous residue I129 in KnChR is surrounded by bulky hydrophobic residues and is positioned proximal to the Schiff-base, at a distance of 4.9 Å (Fig. 5b). The I129Q mutation (KnChortRQ) showed a 40 nm blue-shifted action spectrum (λmax=410 nm) (Fig. 5a, c). I129Q could form a polar interaction with the Schiff base and thereby lower the energy at the ground state, which would form the basis of this blue-shift. We then combined I129Q and A157G to achieve a further blue-shift. However, the KnChR I129Q/A157G mutant shifted only to 430 nm (Fig. 5c), indicating there is no additive effect of the double mutant.

**Fig. 5.**
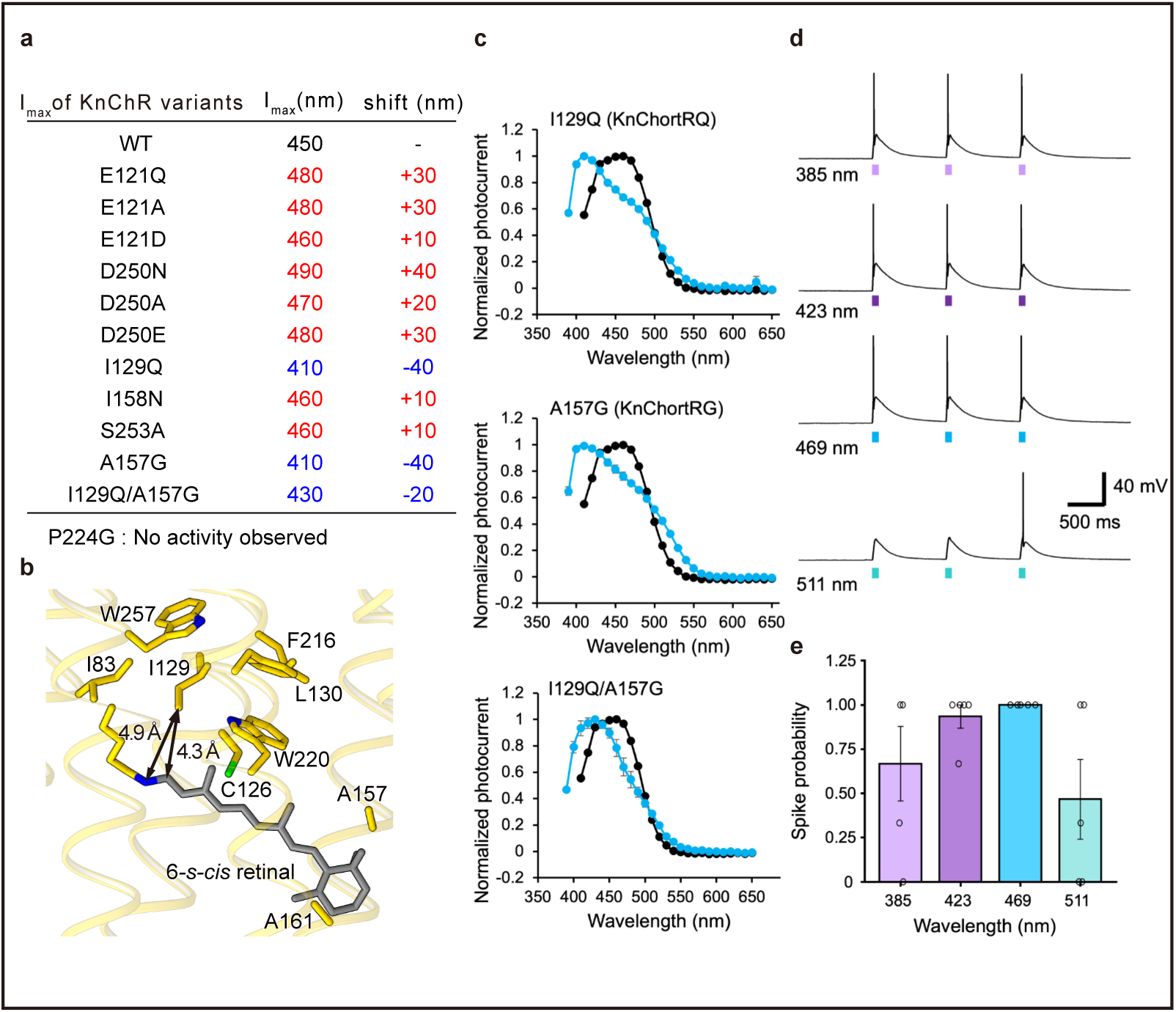
Color-tuning mutants and Optogenetic characterization. **a,** The λ_max_ values of KnChR-272 and KnChR mutants. **b**, Close-up view of I129 and surrounding residues, shown as sticks. **c,** Action spectra of I129Q, A157G, and I129Q/A157G. The wavelength dependency of the photocurrent from KnChR-272 (black) and mutants (blue) is depicted. Membrane voltage was clamped at −60 mV. n = 5 to 6 cells. **d**, Representative responses of a KnChR-272 expressing neuron. The action potentials evoked by 385, 423, 469, and 511 nm wavelengths of light. The 50 ms light pulse was performed as indicated by the colored time points. Light intensities were 1.4 mW/mm^2^. **e**, KnChR-272 expressing neurons were excited by four wavelengths of light at the same light intensities. To compare the success rate of optical stimulation, spike probabilities (total numbers of evoked action potentials / total numbers of light stimulation) are compared. n = 5 cells.

We finally tested the optogenetics applicability of KnChR, particularly with short-wavelength light. Precultured cortical neurons were transfected with KnChR, and optical stimulations were performed. As shown in Fig. 5d, 50 ms light-pulse successfully triggered action potentials. As expected from the action spectrum of KnChR, 423 and 469 nm light are most effective, exhibiting ∼90-100% of spike probability, while less probability was observed with 511 nm light (50%) (Fig. 5e). Notably, 385 nm light was able to excite action potentials with relatively good efficiency (70%). These results indicate that KnChR could be used as an optogenetics tool, particularly with UV or deep-blue light.

## Discussion

In this study, we performed structural and functional analyses of the blue-shifted channelrhodopsin KnChR. Compared to typical channels and GPCR-G-protein complexes^28^, the cryo-EM structural analysis of KnChR was challenging, because KnChR has fewer features such as large domains in the outer membrane region. We succeeded in the structural determination by using nanodiscs^29^. The distinctive C-terminal region covers the intracellular loops of the protein, implying the potential allosteric modulation of conformational changes upon light irradiation. Furthermore, our findings revealed that the 6-*s-cis* retinal configuration serves as a key determinant of blue-shifted excitation. Employing a structure-guided engineering approach, we created the KnChR mutants with the most pronounced blue-shifted excitation (KnChortRG and KnChortRQ), thus harboring the potential for efficacious dual-light utilization in optogenetics. This engineering strategy holds promise for applications to other ChRs, thereby extending the horizons of optogenetic utility.

For cryo-EM studies, nanodiscs are frequently used for the structural determination of ion channels and transporters. In general, nanodisc reconstitution diminishes the unwanted densities of lipids and detergents, while concurrently improving the protein stability. This enhancement facilitates particle alignment during the cryo-EM data processing. This methodology has also been effectively utilized for determining the structures of relatively small transporters. For microbial rhodopsins, nanodisc reconstitution has been used in the cryo-EM structural analyses of the ChRmine trimer^30^ and the Kin4B8 pentamer^31^. We adopted this approach for KnChR, and ultimately determined the cryo-EM structure of the nanodisc-reconstituted KnChR. It is noteworthy that the molecular weight of the KnChR dimer, as extrapolated from the structural model, is 60.8 kDa, underscoring its status as one of the smallest membrane proteins for which cryo-EM structures have been determined. We initially evaluated the nanodisc reconstitution efficiency on a small scale with various membrane scaffold proteins (MSPs) (Supplementary Fig. 2a). The results showed that MSP2N2 is the best for the reconstruction, as it creates nanodiscs with a larger diameter than those of MSP1E3, MSP1E3D1, and MSP1D1^32^. Since MSP1E3D1 was previously employed in the structural analysis of the ChRmine trimer, it is somewhat remarkable that MSP2N2, boasting a larger diameter, successfully facilitated the structure determination of the KnChR dimer. Accordingly, in the structural analyses of moderately-sized membrane proteins, the selection of the appropriate type of MSP warrants careful consideration.

X-ray crystallography has been the pioneering method for the structural elucidations of microbial rhodopsins since the late 1990s^33^. Subsequently, the architectures of GPR^31^, ChRmine^30^, and Kin4B8^31^ have been determined by cryo-EM, a technique that obviates the necessity for crystallization. Notably, the success of this structural analysis of KnChR indicates that the structural examination of all microbial rhodopsins has become fundamentally feasible, given their propensity to form dimers or additional multimeric configurations. Specifically, several X-ray crystal structures of dimeric ChRs used in optogenetics have reported^34,35^, yet a plethora of structures remain unsolved, encompassing optogenetically useful variants, disparate subtypes within a family, and ChRs engineered by machine learning. The acquisition of these structural data would substantively aid in deciphering and engineering the multifaceted diversity of ChRs.

## Acknowledgements

We thank K. Ogomori, C. Harada, and Y. Kanazawa for technical assistance. This work was supported by grants from the Institute for Fermentation Osaka (W.S.), JSPS KAKENHI (grants 19H05777 to W.S., 21H04969 to H.K., 18K06109 to S.P.T., 20K15900 to S.H., and 21H05037 to O.N.), a JST CREST grant (JPMJCR1753 to H.K.), and a JST PRESTO grant (JPMJPR1688 to S.P.T.), the Platform Project for Supporting Drug Discovery and Life Science Research (Basis for Supporting Innovative Drug Discovery and Life Science Research) from the Japan Agency for Medical Research and Development (AMED) under grant number JP22ama121002 (support number 3272 to O.N.) and JP22ama121012 (support number 4893 to H.K.).

## Author contributions

Y.Z.W. performed the structural analysis with assistance from T.T., and H.A. and F.K.S. assisted with the single particle analysis. K.N., R.T., S.P.T., and H.K. performed the electrophysiological characterization of the mutant proteins. W.S. made the expression vector for the structural analysis and designed the experiment. The manuscript was mainly prepared by Y.Z.W. and W.S., with assistance from O.N. W.S., H.K., and O.N. supervised the research.

## Methods

### Expression of KnChR

The KnChR gene was subcloned into a pFastBac vector, modified to include a C-terminal EGFP-His tag, followed by a tobacco etch virus (TEV) protease recognition site. The construct was expressed in *sf9* cells using the pFastBac baculovirus system. The baculovirus was used to infect the *sf9* cells when the cells were grown in suspension to a density of 3.5 × 10^6^ cells mL^-1^, and the cells were then cultured at 27°C for 42 hours. All*-trans*-retinal was supplemented to a final concentration of 10 µM in the cell medium, 24 hours after the infection.

### Purification of KnChR with GDN

The cells were harvested by centrifugation at 5,000 g for 12 minutes. The pellets were first disrupted in solubilization buffer, containing 150 mM NaCl, 20 mM Tris-HCl, pH 8.0, 10% glycerol, 1 mM aprotinin, 1 mM leupeptin, 1 mM pepstatin, 1 mM phenylmethylsulfonyl fluoride (PMSF), 1% n-Dodecyl-β-D-maltoside (DDM) (Calbiochem), and 0.2% cholesteryl hemisuccinate (CHS), using a glass Dounce homogenizer. Both the supernatant and insoluble fractions were then collected and solubilized in the same solubilization buffer for 30 minutes at 4°C. The supernatant was separated from insoluble material by ultracentrifugation at 15,000*g* for 20 minutes, and incubated with TALON resin (Clontech) for 30Cminutes. The resin was washed with ten column volumes of buffer, containing 20 mM Tris-HCl, pHC8.0, 500 mM NaCl, 0.05% glyco-diosgenin (GDN) (Anatrace), and 15 mM imidazole. The receptor was eluted in buffer, containing 20 mM Tris-HCl, pHC8.0, 500 mM NaCl, 0.05% GDN, and 200 mM imidazole. The flow-through fraction was collected and concentrated using an Amicon Ultra 10 kDa molecular weight cutoff centrifugal filter unit (Merck Millipore). The concentrated sample was then subjected to size-exclusion chromatography on a Superose™ 6 Increase 10/300 GL column (GE Healthcare Life Sciences), in buffer containing 150 mM NaCl, 20 mM Tris-HCl, pHC8.0, and 0.01% GDN. Peak fractions were pooled and frozen in liquid nitrogen (Supplementary Fig. 1b).

### Purification of KnChR with LMNG

We performed detergent screening to enhance the purification efficiency, and found that Lauryl Maltose Neopentyl Glycol (LMNG) is best for KnChR solubilization. After the cells were harvested, the pellets were disrupted in solubilization buffer containing 150 mM NaCl, 20 mM Tris-HCl, pH 8.0, 10% glycerol, 1 mM aprotinin, 1 mM leupeptin, 1 mM pepstatin, 1 mM PMSF, 1% Lauryl Maltose Neopentyl Glycol (LMNG), and 0.2% CHS, using a glass Dounce homogenizer. Both the supernatant and insoluble fractions were then collected and solubilized in the same solubilization buffer for 2 hours at 4°C. The supernatant was separated from insoluble material by ultracentrifugation at 15,000*g* for 20 minutes, and incubated with GFP nanobody-coupled resin at 4°C for 2 hours. To cleave the tags, 1 mg mL^-1^ TEV protease was added and the sample was rotated for 12 hours at 4°C. The collected elution fraction was reloaded on the Ni-NTA Superflow resin (QIAGEN), with 20 mM imidazole, to trap the cleaved GFP-His tags and the overloaded TEV protease. The flow-through fraction was collected, concentrated, and then loaded onto a Superose™ 6 Increase 10/300 GL column (GE Healthcare Life Sciences), in buffer containing 150 mM NaCl, 20 mM Tris-HCl, pH 8.0, and 0.01% LMNG. Protein-containing fractions were pooled and frozen in liquid nitrogen.

### Nanodisc reconstitution

To determine whether KnChR can be reconstituted in nanodiscs, the protein in LMNG micelles was mixed with a film of lipid SoyPC (initially dissolved in chloroform) resuspended with 1% cholic acid and several MSPs, at a molar ratio of 1:200:4, respectively, and incubated at 4°C for 1 hour. The LMNG detergent was removed by adding Bio-Beads SM2 (Bio-Rad) to 40 mg mL^-1^, followed by gentle agitation. The Bio-Beads were replaced with fresh ones after 2 hours, and this second batch (equal amount) was incubated overnight at 4°C. The Bio-Beads were then removed, and the solution was ultracentrifuged before size-exclusion chromatography. The ultracentrifuged sample was purified by size-exclusion chromatography on a Superose™ 6 Increase 10/300 GL column (GE Healthcare Life Sciences), equilibrated with buffer containing 20 mM Tris-HCl, pH 8.0, and 150 mM NaCl (Supplementary Fig. 2a).

For the cryo-EM grid preparation of the KnChR, the protein was reconstituted in nanodiscs composed of SoyPC and 2N2, as described above. After the Bio-Beads were removed, the solution was ultracentrifuged and then purified by size-exclusion chromatography. The peak fractions of the protein were collected and concentrated to 4 mg mL^-1^, using a centrifugal filter unit (10 kDa molecular weight cut-off; Merck Millipore).

### Sample vitrification

A 3 µl portion of the sample was loaded onto glow-discharged holey carbon grids (Quantifoil Cu/Rh 300 mesh R1.2/1.3), which were then plunge-frozen in liquid ethane, using a Vitrobot Mark IV (Thermo Fischer Scientific).

For KnChR purified in GDN, data collections were performed on a 200kV Titan Talos G3i microscope (Thermo Fisher Scientific) equipped with a BioQuantum K2 imaging filter and a Falcon 3EC direct electron detector (Gatan), using the EPU software (Thermo Fisher’s single-particle data collection software). Images were obtained at a dose rate of about 71.939 e Å^-2^, with a defocus range from −0.8 to −1.8 μm. The total exposure time was 4.25 seconds, with 50 frames recorded per micrograph. A total of 5,022 videos were collected. All acquired movies in super-resolution mode were binned by 2, dose-fractionated, and subjected to beam-induced motion correction implemented in cryoSPARC v3.3^36^.

For KnChR reconstituted in nanodiscs, images were obtained at a dose rate of about 49.26 e Å^-2^, with a defocus range from −0.6 to −1.6 μm. The total exposure time was 2.11Cseconds, with 48 frames recorded per micrograph. A total of 8,554 videos were collected. All acquired movies in super-resolution mode were binned by 2, dose-fractionated, and subjected to beam-induced motion correction implemented in cryoSPARC v3.3^36^.

### cryo-EM single particle analysis

The contrast transfer function (CTF) parameters were estimated by using patch CTF estimation in cryoSPARC v3.3^36^. Particles were initially picked from a small fraction with Gaussian blob picking and subjected to 2D classification.

For KnChR purified in GDN, class averages showing reasonable features of the KnChR dimer in various orientations were selected as templates, and particles were then picked using the templates. Particles from these class averages generated an ab initio model in cryoSPARC. For each full dataset, extracted particles were down-sampled to 3.32 Å, followed by two rounds of 2D classification to remove ‘junk’ particles. Non-uniform refinement with the ab initio model as a reference was performed using C2 symmetry. Finally, the 82,753 particles in the best class were reconstructed using non-uniform refinement, resulting in a 3.45 Å resolution reconstruction, with the gold-standard Fourier shell correlation (FSCC=C0.143) criteria. The processing strategy is described in Supplementary Fig. 1c.

For KnChR reconstituted in nanodiscs, class averages showing reasonable features of the KnChR dimer in various orientations were selected as templates for the training of the Topaz particle picking program^37^ and particles were then picked accordingly. Particles from these class averages were used to generate an ab initio model in cryoSPARC. For each full dataset, extracted particles were down-sampled to 3.32CÅ, followed by two rounds of 2D classification to remove ‘junk’ particles. Heterogenous refinement into 3 classes with the ab initio model as a reference was performed using C2 symmetry. After multiple rounds of heterogenous refinement, particles were re-extracted with a pixel size of 1.25 Å and a box size of 180 pixels. Multiple rounds of local CTF refinement and non-uniform refinement were performed using cryoSPARC. Finally, the 87,244 particles in the best class were reconstructed using non-uniform refinement, resulting in a 3.01 Å resolution reconstruction, with the gold-standard Fourier shell correlation (FSCC=C0.143) criteria. The processing strategy is described in Supplementary Fig. 2b.

### Model building and refinement

The quality of the map with the nanodisc-reconstituted protein was sufficient to build a model manually in Coot. The model building was facilitated by the AlphaFold-predicted KnChR model. We manually readjusted the model into the density map using Coot and refined it using phenix.real_space_refine (v.1.19)^38,39^. The final model of KnChR contained residues 27-291, all-*trans-*retinal, and 24 water molecules. Detailed parameters are listed in Table 1. Structural analysis and figure preparations were performed with the USCF ChimeraX (V1.6.1)^40^ and CueMol (Ver2.2.3.443) software.

### Expression plasmid

The pKnChR-eYFP plasmid for expression in mammalian cells was described previously^20^. Site-directed mutagenesis was performed using a QuikChange site-directed mutagenesis kit (Agilent, CA, USA) or KOD -Plus- Mutagenesis Kit (Toyobo, Osaka, Japan). All constructs were verified by DNA sequencing (Fasmac Co., Ltd., Kanagawa, Japan).

### Virus preparation

The AAV7m8 KnChR-Venus vector was produced by VectorBuilder (Yokohama, Japan).

### Cell culture

ND7/23 cells, a hybrid cell line derived from neonatal rat dorsal root ganglia neurons fused with mouse neuroblastoma cells, were grown on collagen-coated coverslips in Dulbecco’s modified Eagle’s medium (Fujifilm Wako Pure Chemical Corporation, Osaka, Japan), supplemented with 2.5 μM all-*trans* retinal and 5% fetal bovine serum, under a 5% CO_2_ atmosphere at 37°C. The expression plasmids were transiently transfected by using FuGENE HD (Promega, Madison, WI, USA), according to the manufacturer’s instructions. Electrophysiological recordings were then conducted 16–36 h after the transfection. Successfully transfected cells were microscopically identified by eYFP fluorescence prior to the measurements.

Cortical neurons were isolated from embryonic day 16 Wistar rats (Japan SLC, Inc., Shizuoka, Japan) using Nerve-Cells Dispersion Solutions (Fujifilm Wako Pure Chemical Corporation) according to the manufacturer’s instructions, and grown in the neuron culture medium (FUJIFILM Wako Pure Chemical Corporation) under a 5% CO_2_atmosphere at 37°C. Cultured cortical neurons were infected at day in vitro (DIV) 7, using an adeno-associated virus (1×10^10^ GC/mL). Viral dilutions (1 μL) were added to cultured cortical neurons seeded on coverslips in 24-well plates, which were incubated at 37°C. At DIV21, electrophysiological recordings were conducted using neurons identified by fluorescence under a conventional epifluorescence system.

### Electrophysiology

All experiments were performed at room temperature (23±2°C). For whole-cell voltage clamp recording, photocurrents were recorded by using an Axopatch 200B amplifier (Molecular Devices, Sunnyvale, CA, USA). Data were filtered at 5 kHz, sampled at 10 kHz, digitized by Digidata1550 (Molecular Devices, Sunnyvale, CA, USA), and stored in a computer. The pipette resistance was between 3–6 MΩ. The internal pipette solution for whole-cell voltage-clamp recording contained (in mM) 120 NaCl, 1 KCl, 2 CaCl_2_, 2 MgCl_2_, 5 EGTA, and 25 HEPES, adjusted to pH 7.2. The extracellular solution contained (in mM) 140 NaCl, 1 KCl, 2 CaCl_2_, 2 MgCl_2_, and 10 HEPES, adjusted to pH 7.2. For recording action spectra, the pipette solution contained (in mM) 126 Na aspartate, 0.5 CaCl_2_, 2 MgCl_2_, 5 EGTA, 25 HEPES, and 12.2 N-methyl D-glucamine, adjusted to pH 7.4 with citric acid. The standard extracellular solution contained (in mM) 150 NaCl, 1.8 CaCl_2_, 1 MgCl_2_, 10 HEPES, 10 N-methyl D-glucamine, and 5 glucose, adjusted to pH 7.4 with N-methyl D-glucamine. Irradiation at 470 nm was performed with a collimated LED (LCS-0470-03-22, Mightex, Toronto, Canada) controlled by computer software (pCLAMP10.7, Molecular Devices, Sunnyvale, CA, USA). Light power was measured directly through the objective lens of a microscope by a power meter (LP1, Sanwa Electric Instruments Co., Ltd., Tokyo, Japan). All action spectra were measured at the same light intensity in the range of 390 nm to 650 nm by an OSG xenon light source (Hamamatsu Photonics, Hamamatsu, Japan).

For whole-cell current clamp recording, action potentials were recorded by using an amplifier IPA (Sutter Instrument, Novato, CA, USA) under a whole-cell patch clamp configuration. Data were filtered at 5 kHz, sampled at 10 kHz, and stored in a computer (SutterPatch, Sutter Instrument, Novato, CA, USA). The pipette resistance was between 5–10 MΩ. The internal pipette solution contained (in mM) 125 K-gluconate, 10 NaCl, 0.2 EGTA, 10 HEPES, 1 MgCl_2_, 3 MgATP, 0.3 Na_2_GTP, 10 Na_2_-phosphocreatine, and 0.1 leupeptin, adjusted to pH 7.4 with KOH. The extracellular Tyrode’s solution contained (in mM) 138 NaCl, 3 KCl, 10 HEPES, 4 NaOH, 2 CaCl_2_, 1 MgCl_2_, and 11 glucose, adjusted to pH 7.4 with KOH. In all cortical neuron experiments, Tyrode’s solution contained 20 μM 6,7-dinitroquinoxaline-2,3-dione (DNQX, Tocris Bioscience, Ellisville, MO, USA), 25 μM d-(−)-2-amino-5-phosphonovaleric acid (D-AP5, Tocris), and 100 μM picrotoxin (Nacalai Tesque, Inc., Kyoto, Japan). The liquid junction potential was 16.3 mV and was compensated. Irradiations at 385, 423, 469 and 511 nm were performed by using a Colibri7 light source (Carl Zeiss, Oberkochen, Germany) controlled by computer software (SutterPatch, Sutter Instrument, Novato, CA, USA). Light power was directly measured through the objective lens of the microscope by a visible light-sensing thermopile (MIR-100Q, SSC Inc., Mie, Japan).

All data in the text and figures are expressed as mean ± SEM and were evaluated with the Mann–Whitney U test for statistical significance, unless otherwise noted. Data were judged as statistically insignificant when P>0.05.

## Data availability

The cryo-EM density map and atomic coordinates for KnChR have been deposited in the Electron Microscopy Data Bank and the PDB, under accession codes XXXX and ZZZZ, respectively.

## References

1. Nagel, G. et al. Channelrhodopsin-2, a directly light-gated cation-selective membrane channel. Proc Natl Acad Sci U S A 100, 13940–13945 (2003).

2. Ernst, O. P. et al. Microbial and Animal Rhodopsins: Structures, Functions, and Molecular Mechanisms. Chem Rev 114, 126–163 (2014).

3. Boyden, E. S., Zhang, F., Bamberg, E., Nagel, G. & Deisseroth, K. Millisecond-timescale, genetically targeted optical control of neural activity. Nat Neurosci 8, 1263– 1268 (2005).

4. Volkov, O. et al. Structural insights into ion conduction by channelrhodopsin 2. Science 358, eaan8862 (2017).

5. Mattis, J. et al. Principles for applying optogenetic tools derived from direct comparative analysis of microbial opsins. Nat Methods 9, 159–172 (2012).

6. Klapoetke, N. C. et al. Independent Optical Excitation of Distinct Neural Populations. Nat Methods 11, 338–346 (2014).

7. Bauer, J. et al. Limited functional convergence of eye-specific inputs in the retinogeniculate pathway of the mouse. Neuron 109, 2457–2468.e12 (2021).

8. Christoffel, D. J. et al. Input-specific modulation of murine nucleus accumbens differentially regulates hedonic feeding. Nat Commun 12, 2135 (2021).

9. Anisimova, M. et al. Spike-timing-dependent plasticity rewards synchrony rather than causality. Cereb Cortex 33, 23–34 (2022).

10. Hooks, B. M., Lin, J. Y., Guo, C. & Svoboda, K. Dual-channel circuit mapping reveals sensorimotor convergence in the primary motor cortex. J Neurosci 35, 4418– 4426 (2015).

11. Chiu, C. Q. et al. Input-Specific NMDAR-Dependent Potentiation of Dendritic GABAergic Inhibition. Neuron 97, 368–377.e3 (2018).

12. Birdsong, W. T. et al. Synapse-specific opioid modulation of thalamo-cortico-striatal circuits. Elife 8, e45146 (2019).

13. Prasad, A. A. et al. Complementary Roles for Ventral Pallidum Cell Types and Their Projections in Relapse. J Neurosci 40, 880–893 (2020).

14. Xia, S.-H. et al. Cortical and Thalamic Interaction with Amygdala-to-Accumbens Synapses. J Neurosci 40, 7119–7132 (2020).

15. Joffe, M. E. et al. Acute restraint stress redirects prefrontal cortex circuit function through mGlu5 receptor plasticity on somatostatin-expressing interneurons. Neuron 110, 1068–1083.e5 (2022).

16. Rindner, D. J., Proddutur, A. & Lur, G. Cell-type-specific integration of feedforward and feedback synaptic inputs in the posterior parietal cortex. Neuron 110, 3760–3773.e5 (2022).

17. Rindner, D. J. & Lur, G. Practical considerations in an era of multicolor optogenetics. Front Cell Neurosci 17, 1160245 (2023).

18. Marshel, J. H. et al. Cortical layer–specific critical dynamics triggering perception. Science 365, eaaw5202 (2019).

19. Kato, H. E. et al. Atomistic design of microbial opsin-based blue-shifted optogenetics tools. Nat Commun 6, 7177 (2015).

20. Tashiro, R. et al. Specific residues in the cytoplasmic domain modulate photocurrent kinetics of channelrhodopsin from Klebsormidium nitens. Commun Biol 4, 1–10 (2021).

21. Kato, H. E. et al. Crystal structure of the channelrhodopsin light-gated cation channel. Nature 482, 369–374 (2012).

22. Kato, H. E. et al. Structural mechanisms of selectivity and gating in anion channelrhodopsins. Nature 561, 349–354 (2018).

23. Tajima, S. et al. Structural basis for ion selectivity in potassium-selective channelrhodopsins. Cell 186, 4325–4344.e26 (2023).

24. Kato, H. E. et al. Atomistic design of microbial opsin-based blue-shifted optogenetics tools. Nature Communications 6, 7177 (2015).

25. Quantum effects in biology. (Cambridge university press, 2014).

26. Sugiyama, Y. et al. Photocurrent attenuation by a single polar-to-nonpolar point mutation of channelrhodopsin-2. Photochem. Photobiol. Sci. 8, 328–336 (2009).

27. Man, D. et al. Diversification and spectral tuning in marine proteorhodopsins. EMBO J 22, 1725–1731 (2003).

28. Zhou, Q. et al. Common activation mechanism of class A GPCRs. eLife 8, e50279 (2019).

29. Borch, J. & Hamann, T. The nanodisc: a novel tool for membrane protein studies. 390, 805–814 (2009).

30. Kishi, K. E. et al. Structural basis for channel conduction in the pump-like channelrhodopsin ChRmine. Cell 185, 672–689.e23 (2022).

31. Chazan, A. et al. Phototrophy by antenna-containing rhodopsin pumps in aquatic environments. Nature 615, 535–540 (2023).

32. Grinkova, Y. V., Denisov, I. G. & Sligar, S. G. Engineering extended membrane scaffold proteins for self-assembly of soluble nanoscale lipid bilayers. *Protein Engineering*, Design and Selection 23, 843 (2010).

33. Grote, M., Engelhard, M. & Hegemann, P. Of ion pumps, sensors and channels — Perspectives on microbial rhodopsins between science and history. Biochimica et Biophysica Acta (BBA) - Bioenergetics 1837, 533–545 (2014).

34. Kim, Y. S. et al. Crystal structure of the natural anion-conducting channelrhodopsin GtACR1. Nature 561, 343–348 (2018).

35. Kato, H. E. et al. Crystal structure of the channelrhodopsin light-gated cation channel. Nature 482, 369–374 (2012).

36. Punjani, A., Rubinstein, J. L., Fleet, D. J. & Brubaker, M. A. cryoSPARC: algorithms for rapid unsupervised cryo-EM structure determination. Nat Methods 14, 290–296 (2017).

37. Bepler, T. et al. Positive-unlabeled convolutional neural networks for particle picking in cryo-electron micrographs. Nat Methods 16, 1153–1160 (2019).

38. Adams, P. D. et al. PHENIX: a comprehensive Python-based system for macromolecular structure solution. Acta Cryst D 66, 213–221 (2010).

39. Afonine, P. V. et al. Real-space refinement in PHENIX for cryo-EM and crystallography. Acta Cryst D 74, 531–544 (2018).

40. Goddard, T. D. et al. UCSF ChimeraX: Meeting modern challenges in visualization and analysis. Protein Science 27, 14–25 (2018).

