## Supplemental Figures 1-5 for "Cryo-EM structure of a blue-shifted channelrhodopsin from *Klebsormidium nitens*"

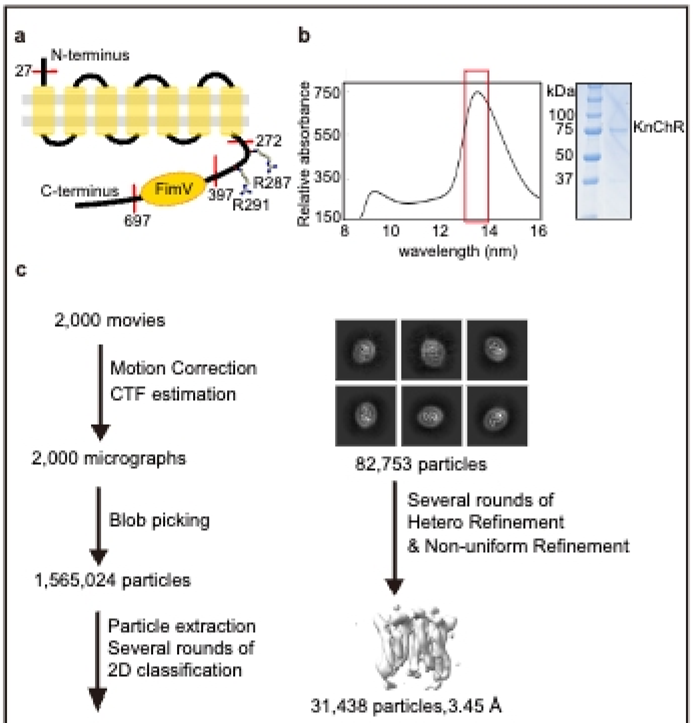


**Supplementary Fig 1: Cryo-EM analysis of KnChR in detergent micelles.** **a**, Schematic representation of KnChR. **b**, Gel-filtration chromatogram and SDS-PAGE of the purified KnChR in LMNG/CHS micelles. **c**, Flow chart of the cryo-EM data processing for the KnChR in LMNG/CHS micelles, including particle projection selection,classification, and 3D density map reconstruction. Details are provided in the Methods section.

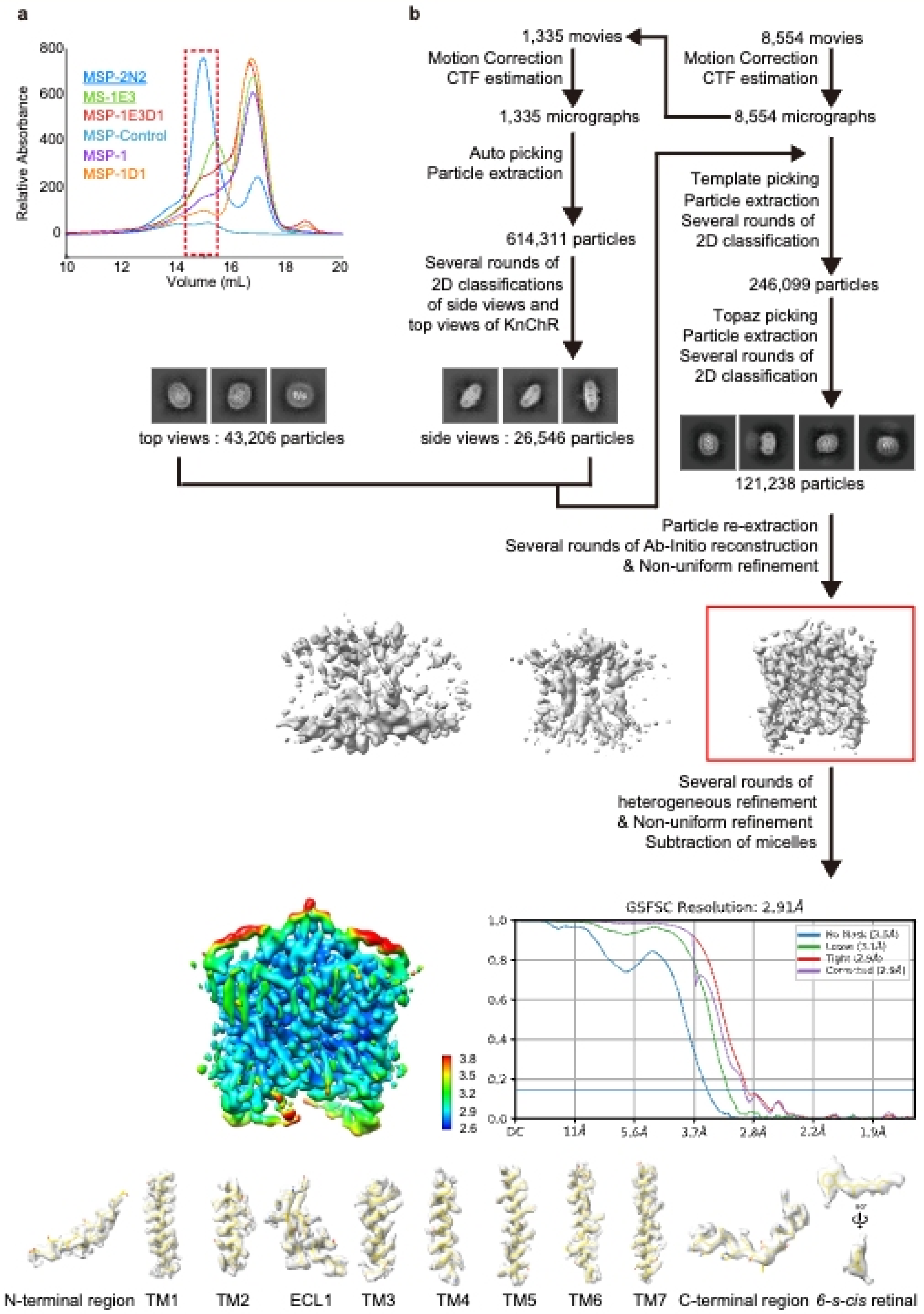


**Supplementary Fig 2: Cryo-EM analysis of KnChR In nanodisc. a**, Evaluation of nanodisc reconstruction KnChR on a small scale. The reconstituted sample was analyzed by Fluorescence-detection size-exclusion chromatography (FSEC) monitoring the tryptophan. **b**, Flow chart of the cryo-EM data processing for the nanodisc-reconstituted KnChR, including particle projection selection, classification, and 3D density map reconstruction, local resolution map, and FSC-curves. The cryo-EM density map and model of the receptor are also shown for all seven transmembrane α-helices, N- and C-terminal regions, and 6-*s-cis* retinal. Details are provided in the Methods section.

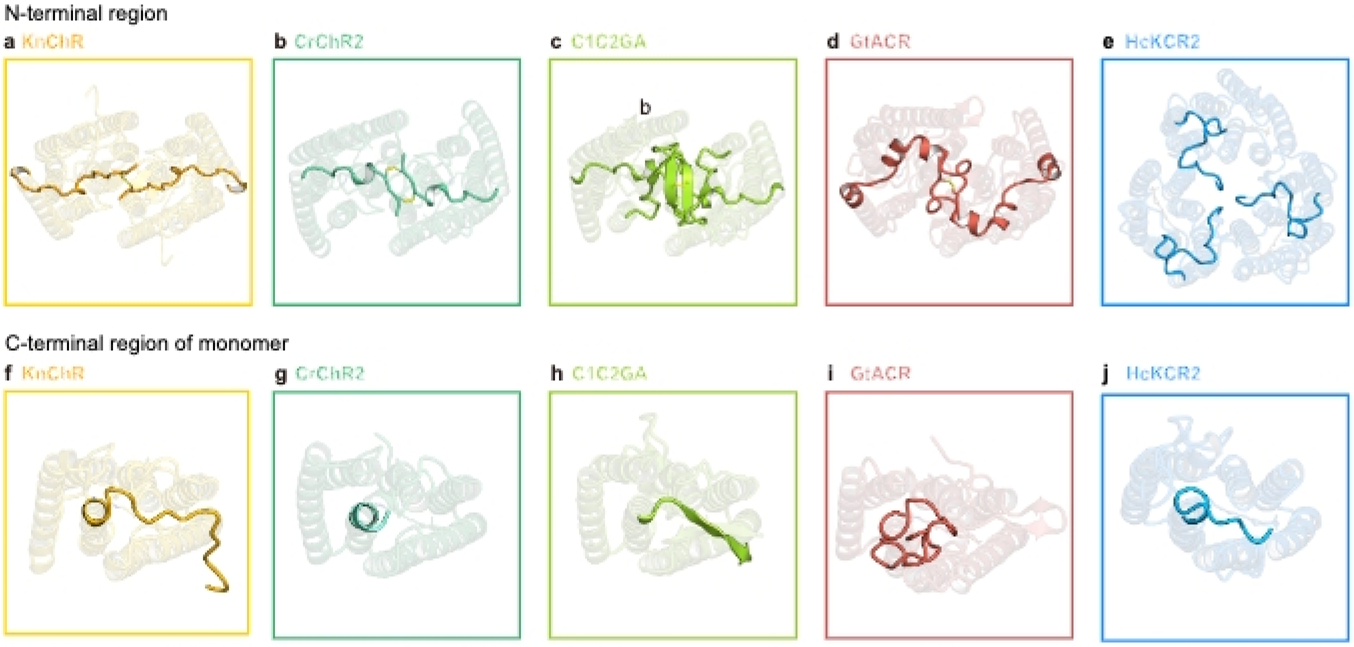


**Supplementary Fig 3: Structural comparison of ChRs. a–e,** Structural comparison of multimeric interfaces of KnChR (**a**), CrChR2 (PDB ID: 6EID) (**b**), C1C2 (PDB ID: 3UG9) (**c**), GtACR1 (PDB ID: 6CSM) (**d**), and HsKCR2 (PDB ID: 8H87) (**e**). **f–j**, Structural comparison of the C-terminal region of KnChR (**f**), CrChR2 (PDB ID: 6EID) (**g**), C1C2 (PDB ID: 3UG9) (**h**), GtACR1 (PDB ID: 6CSM) (**i**), and HsKCR2 (PDB ID: 8H87) (**j**).

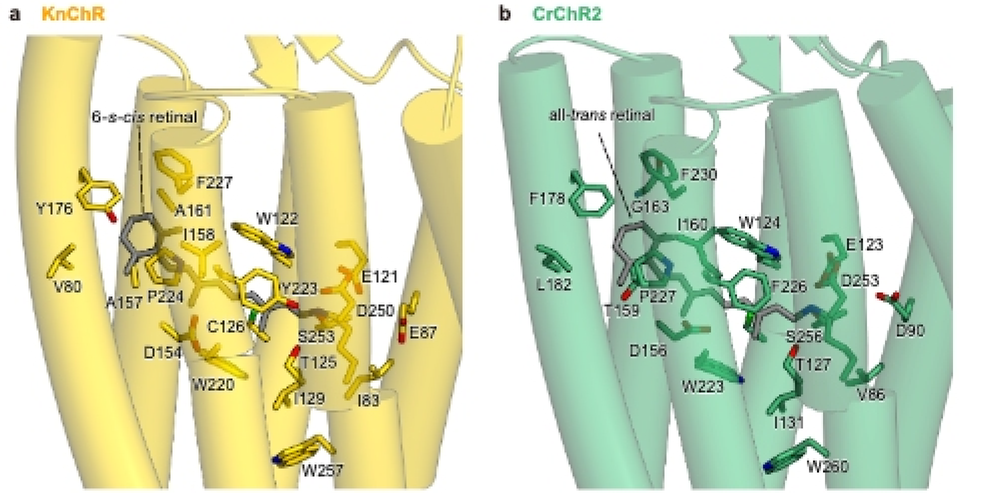


**Supplementary Fig 4: Retinal binding sites in KnChR and CrChR2.a**, The retinal chromophore and the residues within 4.5 Å in KnChR (**a**) and CrChR2 (PDB ID: 6EID) (**b**), involved in retinal binding.

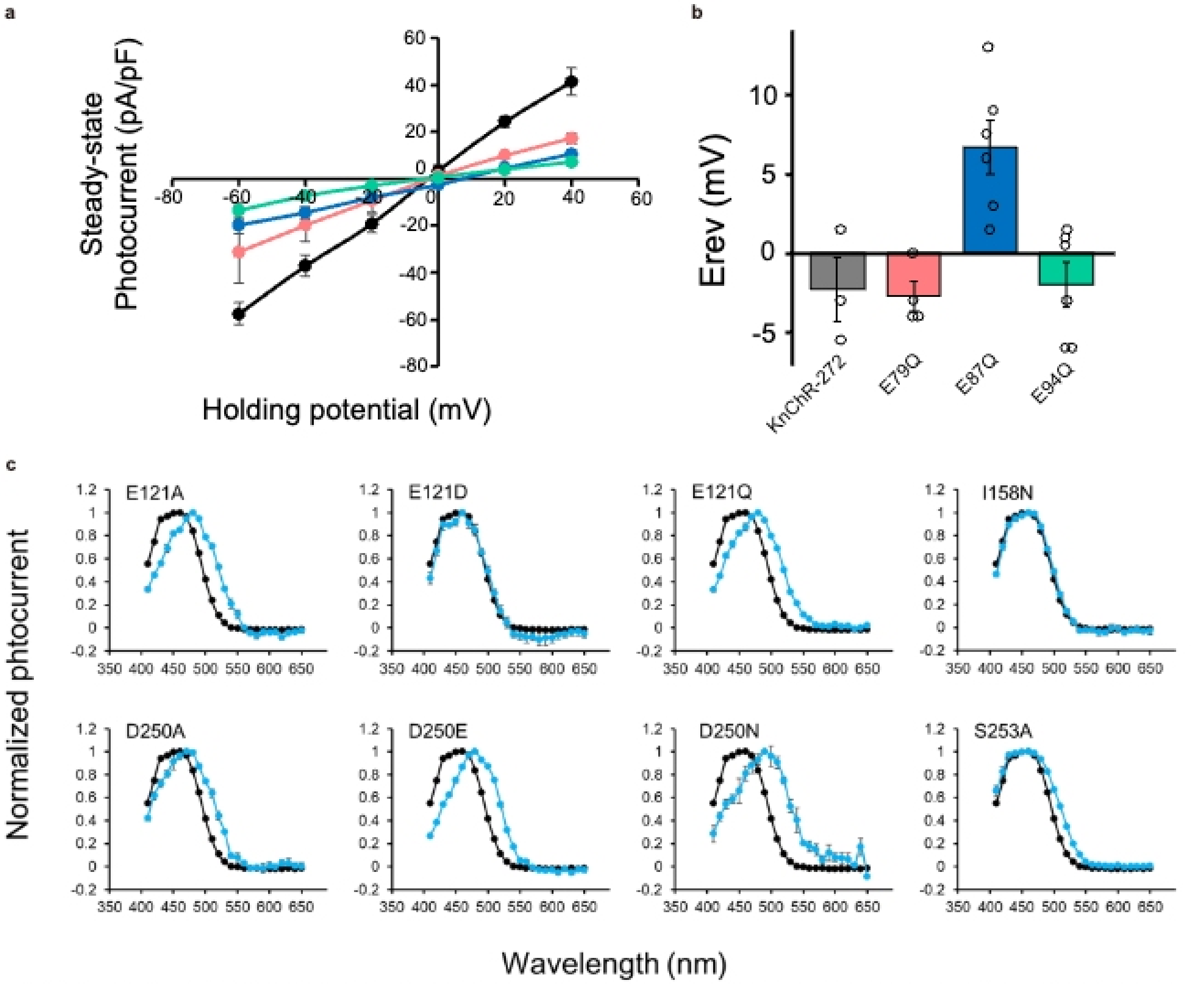


**Supplementary Fig 5: Electrophysiological properties of KnChR and mutants. a**, I-V relationship of the steady-state photocurrents. n = 3 to 6 cells. **b**, Reversal potential (Erev) for KnChR-272 and mutants. n = 3 to 6 cells. **c**, Action spectra of KnChR-272 mutants. Wavelength dependency of the photocurrent from KnChR-272 (black) and mutants (blue) was depicted. Membrane voltage was clamped at -60mV. n = 3 to 7 cells.
